# Neurobiological Signatures of Endometriosis: Characterizing Pain, Cognition, and Brain Morphology

**DOI:** 10.64898/2026.01.05.696004

**Authors:** Elle M. Murata, Gabriella Natividad, Andrea Gabay, Hannah Grotzinger, Tyler Santander, Averi Giudicessi, Morgan Fitzgerald, Allesandra Iadipaolo, Sanjay K. Agarwal, Matthew S. Panizzon, Emily G. Jacobs

**Affiliations:** Department of Psychological & Brain Sciences, University of California, Santa Barbara, CA; Department of Psychiatry, University of California, San Diego, La Jolla, CA; San Diego State University/ University of California, San Diego Joint Doctoral Program in Clinical Psychology, San Diego, California; Department of Neurosciences, University of California, San Diego, La Jolla, California; Department of Psychological and Brain Sciences, Boston University, Boston, MA; Neurosciences Graduate Program, University of California, San Diego, CA; Center for Endometriosis Research and Treatment, Department of Obstetrics, Gynecology and Reproductive Sciences, University of California, San Diego, La Jolla, CA; Center for Behavior Genetics of Aging, University of California, San Diego, La Jolla, CA; Neuroscience Research Institute, University of California, Santa Barbara, CA

**Author notes:** These authors contributed equally to the work (Matthew S. Panizzon, Emily G. Jacobs). **Author Contributions:** The current study was conceived and designed by E.M.M., S.K.A., M.S.P., and E.G.J. Data collection was completed by E.M.M., G.N., A.G., T.S., H.G., A.G., M.F., A.I, and S.K.A. Data analyses were completed by E.M.M. and T.S. The original draft was written by E.M.M. **Competing Interest Statement:** None.

## Abstract

Endometriosis affects approximately 10% of reproductive-age women worldwide and is associated with substantial pain and mental health burden, yet its neurobiological correlates remain poorly characterized. Neuroimaging studies of endometriosis are particularly limited. Here we present a comprehensive, multidimensional investigation of the neurobiology of endometriosis, integrating structural and diffusion neuroimaging, biofluid measures, neuropsychological testing, and detailed health and psychosocial phenotyping in individuals with and without endometriosis. Compared with controls, individuals with endometriosis exhibited higher levels of pain, depression, and anxiety and performed worse across multiple cognitive domains. Global and regional brain morphometric measures did not differ between groups; however, endometriosis-specific pain was associated with altered gray matter volume and white matter microstructure across distributed pain-related neural circuits. These findings identify pain-related neurobiological signatures of endometriosis and establish a foundation for mechanistic and translational studies of its central nervous system effects.

---

Endometriosis is one of the most common reproductive disorders, affecting one in ten women of reproductive age (1). The condition is characterized by the growth of endometrial-like tissue outside the uterus and is associated with debilitating symptoms including chronic pelvic pain, dyspareunia, and infertility. Although highly prevalent, endometriosis remains under-researched and the underlying etiology remains poorly understood.

Traditionally described as an estrogen-dependent gynecological disorder, endometriosis is increasingly recognized as a systemic condition with broad physiological and psychological consequences. Recent studies mapping the immune, metabolic, and chronic pain-related components of endometriosis are leading to an expanded view of the condition beyond the reproductive tract (2). Individuals with endometriosis consistently report lower quality of life (3) and are significantly more likely to experience poor mental health (4,5). A recent systematic review highlights the substantial psychiatric burden in this population, with prevalence rates of depressive symptoms as high as 98% and anxiety symptoms up to 87% (6). Despite unequivocal evidence linking endometriosis to poor mental health outcomes and systemic physiological changes, the underlying neurological sequelae of the condition remain largely unknown.

Initial neuroimaging evidence implicates the central nervous system in endometriosis. Structural MRI studies have reported altered gray matter volume (GMV) in regions involved in pain perception (7,8), while functional MRI studies have identified distinct connectivity patterns linked to pain severity and affective symptoms (9,10). However, these findings are derived from small cohorts and focus predominantly on gray matter volume, limiting reproducibility and sensitivity to detect subtle, distributed neural alterations. Notably, white matter microstructure, which is central to the integrity of large-scale pain and affective networks, has not been examined in endometriosis. Further, a comprehensive neuropsychological characterization of cognition in this population is lacking. Collectively, existing studies suggest central nervous system involvement in endometriosis, yet leave fundamental questions unanswered regarding the broader neural, cognitive, and psychological consequences of the disorder.

To address this gap, here we provide one of the most detailed, comprehensive, multidimensional characterizations of the neurobiology of endometriosis, incorporating mood, pain, cognition, biofluids, and multimodal neuroimaging measures, spanning white matter microstructure to global morphology. Individuals with endometriosis (n = 60) and age-matched healthy controls (n = 60) completed neuropsychological testing, blood draws, and imaging at the University of California, Santa Barbara (UCSB) or the University of California, San Diego (UCSD), enabling a multidimensional characterization of the disorder. Individuals with endometriosis reported greater depression, anxiety, and pain and exhibited reduced performance on several cognitive measures compared to healthy controls. Global and regional measures of GMV, white matter volume, cerebrospinal fluid, cortical thickness, and white matter microstructure were indistinguishable between cases and controls. However, within cases endometriosis-related pain was negatively associated with GMV and positively associated with white matter microstructure throughout pain circuitry. These findings provide novel evidence for a neurobiological basis of endometriosis, linking pain experience to micro and macrostructural alterations in pain circuitry, thus advancing our understanding of the disorder as one with mental health and neurological components.

## Results

### Demographics

Participant demographics are reported in **Table 1** and **Supplementary Table 1**. The current study included 60 highly characterized control participants (M age = 28.12, SD = 3.54) and 60 participants with endometriosis (M age = 29.88, SD = 6.04). With the exception of years of education, there were no differences in demographic variables or endogenous hormone levels between controls and cases (*p* > 0.05 for all, False Discovery Rate [FDR]-corrected). The control group reported more years of education (M = 17.18, SD = 1.46) compared to the case group (M = 15.52, SD = 2.55), or ∼1.5 years on average.

**Table 1.** Demographic Characteristics. FDR-corrected at *q* < 0.05

| Variable | Control (n=60)<br>Mean (SD) \| Range | Case (n=60)<br>Mean (SD) \| Range | p-value |
| --- | --- | --- | --- |
| Age | 28.12 (3.54) \| 24-40 | 29.88 (6.04) \| 19-40 | 0.11 |
| Body Mass Index | 23.61 (4.78) \| 18.79-42.91 | 25.96 (5.92) \| 18.71-43.9 | 0.09 |
| Education (Years) | **17.18 (1.46) \| 15-20** | **15.52 (2.55) \| 12-20** | **0.0003** |
| Mother Education (Years) | 15.73 (2.97) \| 5-20 | 14.71 (2.68) \| 6-20 | 0.11 |
| Race (% White) | 56.67 | 65.67 | 0.08 |
| Ethnicity (% Hispanic or Latino) | 20.00 | 33.33 | 0.28 |
| Estradiol (pg/mL) | 69.63 (77.99) \| 4.29-401 | 81.19 (93.05) \| 0.85-528 | 0.43 |
| Progesterone (ng/mL) | 1.31 (3.96) \| 0.03-22.3 | 1.02 (2.36) \| 0.04-13.3 | 0.51 |
| Testosterone (ng/dL) | 35.36 (21.34) \| 4.65-136 | 31.65 (19.09) \| 2.55-125 | 0.46 |
| Follicle-Stimulating Hormone (mIU/mL) | 5.03 (2.47) \| 0.89-17.28 | 4.93 (5.43) \| 0.2-41.85 | 0.11 |
| Luteinizing Hormone (mIU/mL) | 6.39 (3.61) \| 2.18-19.78 | 7.62 (6.34) \| 0.2-30.03 | 0.74 |

### Cases demonstrated poorer mood and cognitive outcomes

Compared to healthy controls, the endometriosis group displayed worse outcomes on physical functioning, role limitations due to physical health, role limitations due to emotional problems, energy/ fatigue, emotional well-being, pain, general health, and sleep quality (*p* < 0.05 for all, FDR-corrected; **Figure 2B)**. Greater global adverse childhood events, anxiety, depression, post-traumatic stress symptoms, pelvic pain, and chronic pain scores were also reported in this group (*p* < 0.05 for all, FDR-corrected), reflecting established clinical characteristics of endometriosis.

**Figure 1.**
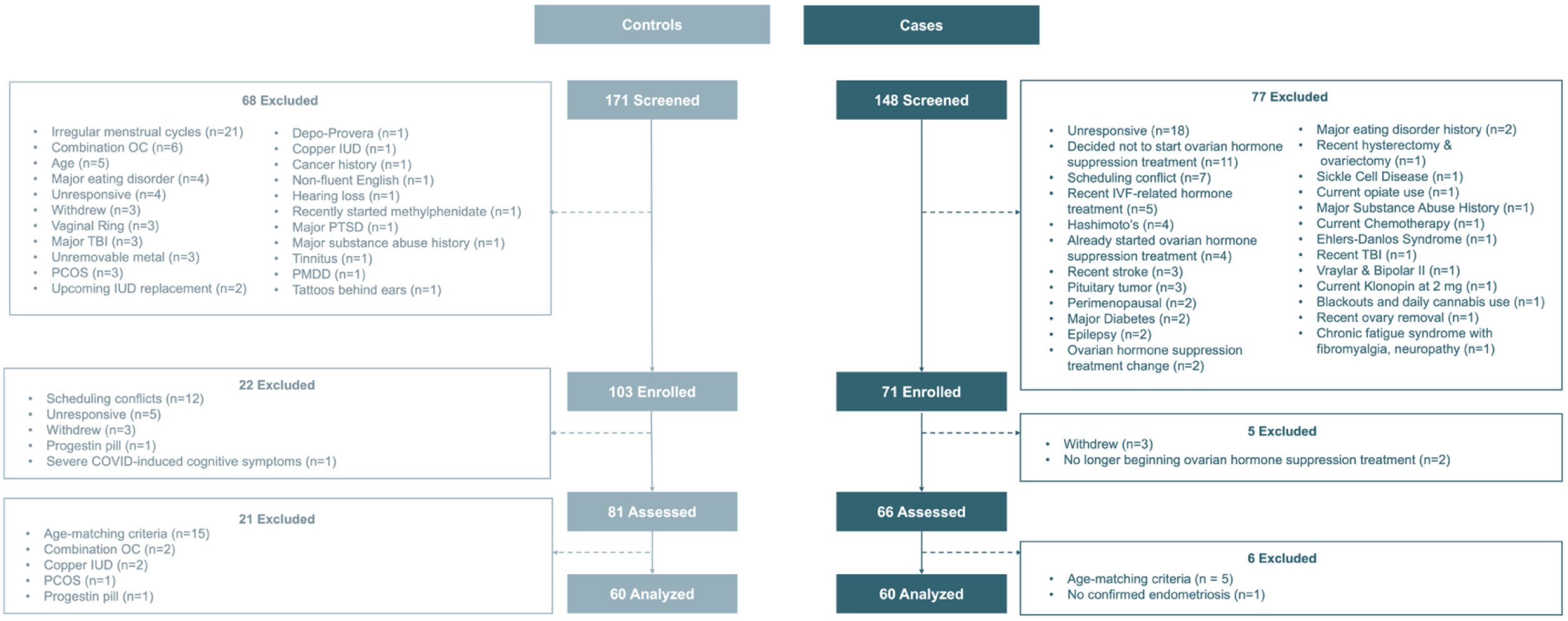
Participant Enrollment. OC = Oral Contraceptive, TBI = Traumatic Brain Injury, PCOS = Polycystic Ovary Syndrome, IUD = Intrauterine Device, PTSD = Post-Traumatic Stress Disorder, PMDD = Premenstrual Dysphoric Disorder, IVF = In-Vitro Fertilization

**Figure 2.**
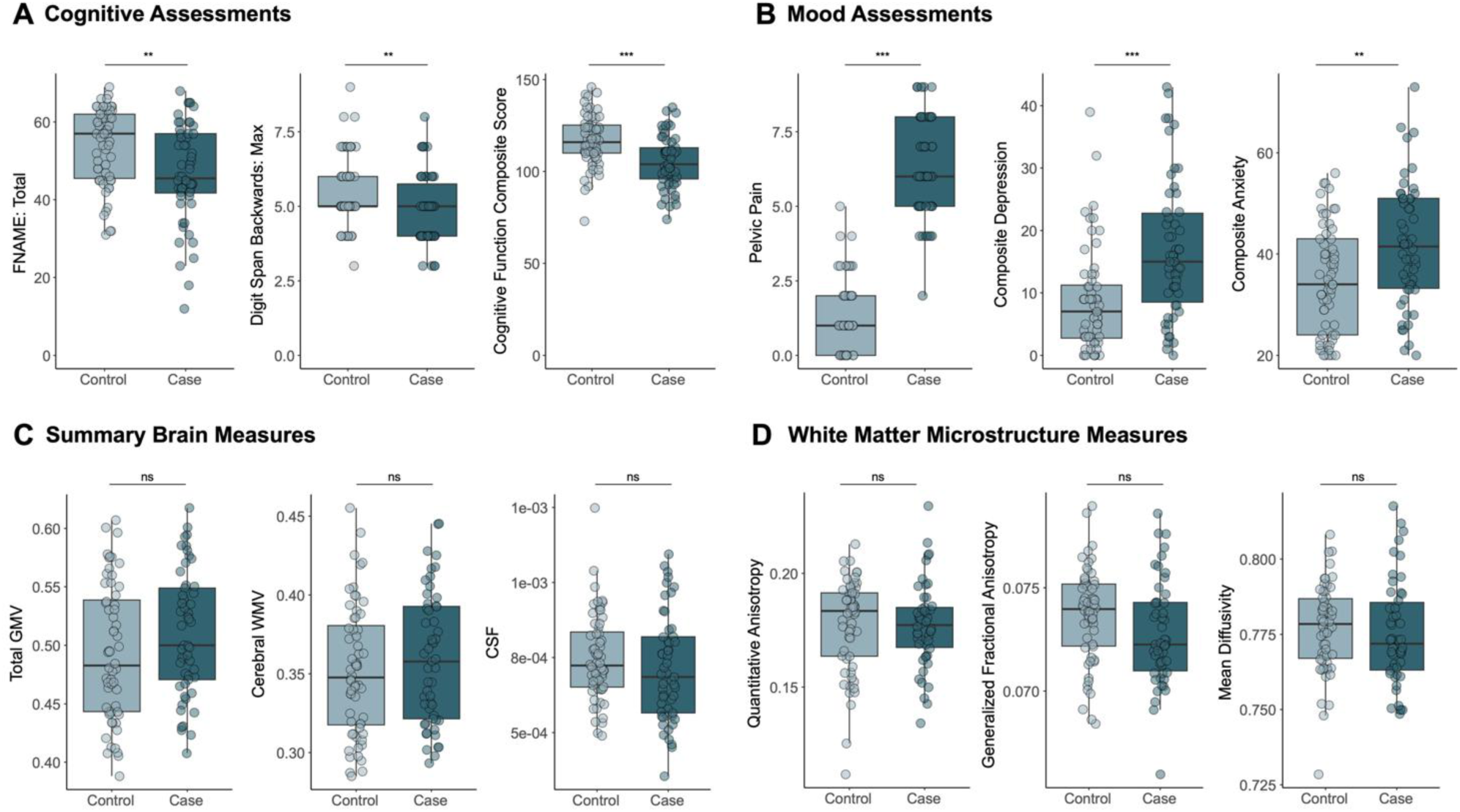
Neurocognitive Measures. All FDR-corrected at *q* < 0.05. FNAME = Face-Name Associative Memory Exam. GMV = Gray Matter Volume, WMV = White Matter Volume, CSF = Cerebrospinal Fluid. \**p*< 0.05, \*\**p*< 0.01, \*\*\**p*< 0.001.

There were no differences between cases and controls on physical activity time logged (*p* > 0.05, FDR-corrected). Full mood and lifestyle results are reported in **Supplementary Table 2**.

After controlling for baseline differences in education, multivariate linear regression models demonstrated that being in the case group was a significant negative indicator of performance on Face-Name Associative Memory Exam (ß = −6.18, SE = 2.27, *p* = 0.03), Stroop test (ß = −5.09, SE = 2.21, *p* = 0.049), Selective Reminding Verbal Memory Consistent Long-Term Retrieval (ß = −6.10, SE = 2.60, *p* = 0.049), and Digit Span Backwards (ß = −0.59, SE = 0.24, *p* = 0.046) **(Figure 2A**). Being in the case group was a significant negative predictor of processing speed on the NIH Toolbox Pattern Comparison test (ß = −5.10, SE = 1.82, *p* = 0.036). Additionally, age-corrected standard scores from the NIH Toolbox battery were higher in the control group compared to the case group (crystallized cognition composite: ß = −7.31, SE = 2.82, *p* = 0.042; overall cognitive function composite: ß = −8.14, SE = 2.67, *p* = 0.034). Full results are reported in **Supplementary Tables 3-4**.

When either the composite depression score or endometriosis-specific pain (as assessed by the Endometriosis Health Profile pain subscale) were included as a covariate in regression models predicting cognitive performance, the independent effect of group status was attenuated. However, depression and endometriosis-specific pain did not correlate with cognitive outcomes within the endometriosis group (*p* > 0.05, FDR-corrected), suggesting that depression or pain and cognitive difficulties represent co-occurring manifestations of endometriosis, rather than serving as direct mechanisms of cognitive performance.

### Global brain metrics reveal no uniform differences between controls and cases

Seven participants did not complete a structural T1-weighted scan, resulting in 113 participants included in the final brain morphology analyses. Diffusion imaging data were available for 108 participants. There were no significant group differences in GMV, cortical thickness, subcortical morphology, or white matter microstructure (all *p* > 0.05) (**Figure 2C-D**). Full results are presented in **Supplementary Tables 5-7**.

### Within cases, endometriosis-related pain is associated with alterations in brain morphology

Within the endometriosis group, Pearson’s two-tailed correlations (FDR-corrected) revealed significant negative associations between endometriosis-related pain and GMV in several cortical regions: right rostral anterior cingulate: *r* (47) = −0.40, *p* = 0.01; right inferior temporal: *r* (47) = −0.40, *p* = 0.01; left entorhinal: *r* (47) = −0.37, *p* = 0.02; left inferior temporal: *r* (47) = - 0.43, *p* = 0.007; left middle temporal: *r* (47) = −0.40, *p* = 0.01; left posterior cingulate: *r* (47) = - 0.34, *p* = 0.03 (**Figure 3)**, in addition to the right middle temporal, right lateral orbitofrontal, and left banks superior temporal sulcus regions (*p* < 0.05 for all). Partial correlations including age as a covariate yielded comparable results. Greater reported total pelvic pain was also negatively correlated with GMV in overlapping regions in cases (*p* < 0.05 for all, FDR-corrected), but not in controls (*p* > 0.05 for all).

**Figure 3.**
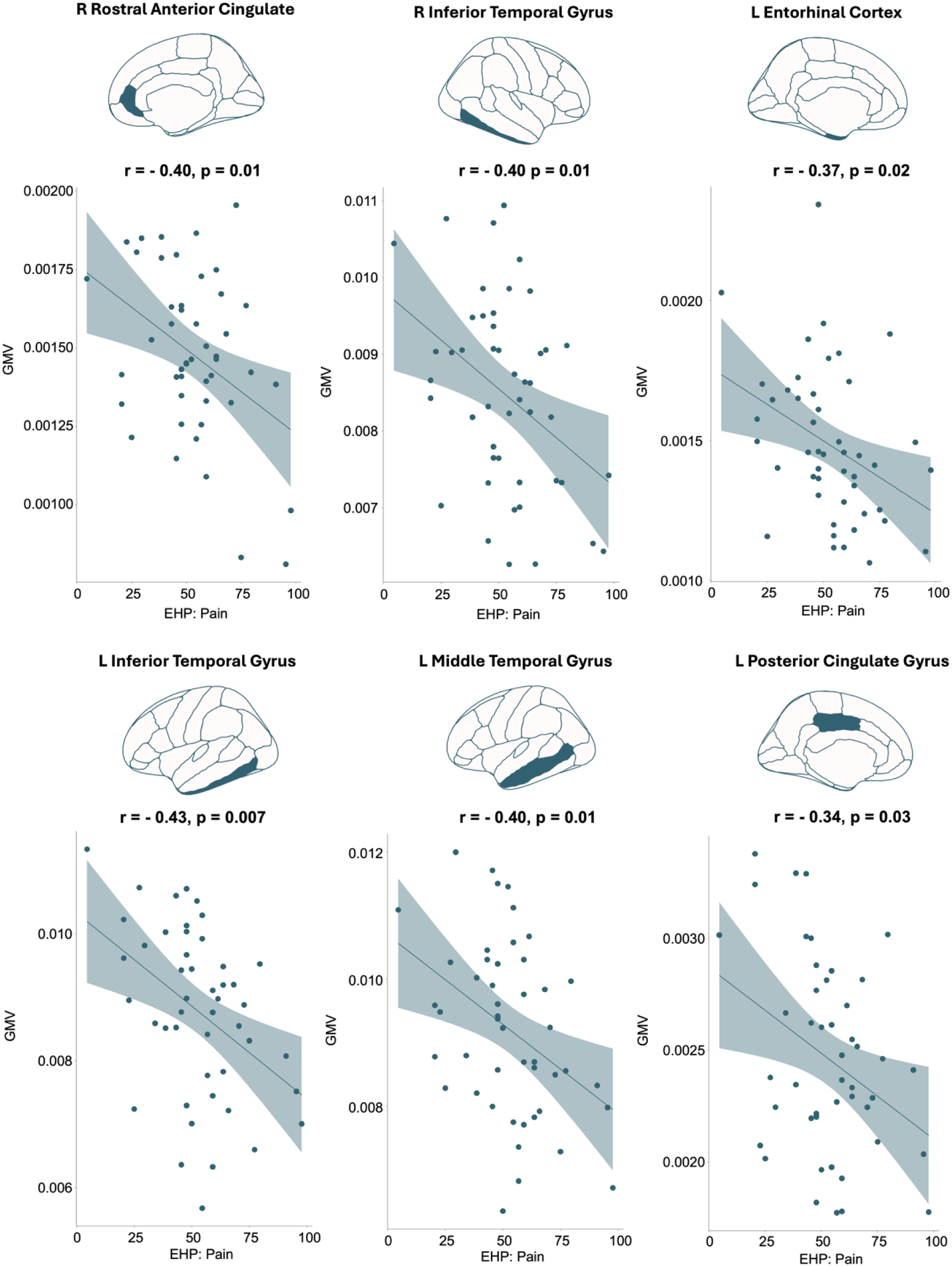
Endometriosis-related pain is negatively associated with GMV. All FDR-corrected at *q* < 0.05. EHP = Endometriosis Health Profile, GMV = Gray Matter Volume.

Several white matter tracts demonstrated a positive relationship between quantitative anisotropy and endometriosis-related pain (FDR, *q* < 0.05, **Figure 4**), including the corpus callosum, cingulum bundle, corticopontine, corticospinal, and superior longitudinal fasciculus tracts. See **Supplementary Table 8** for the full list of quantitative anisotropy tract results.

**Figure 4.**
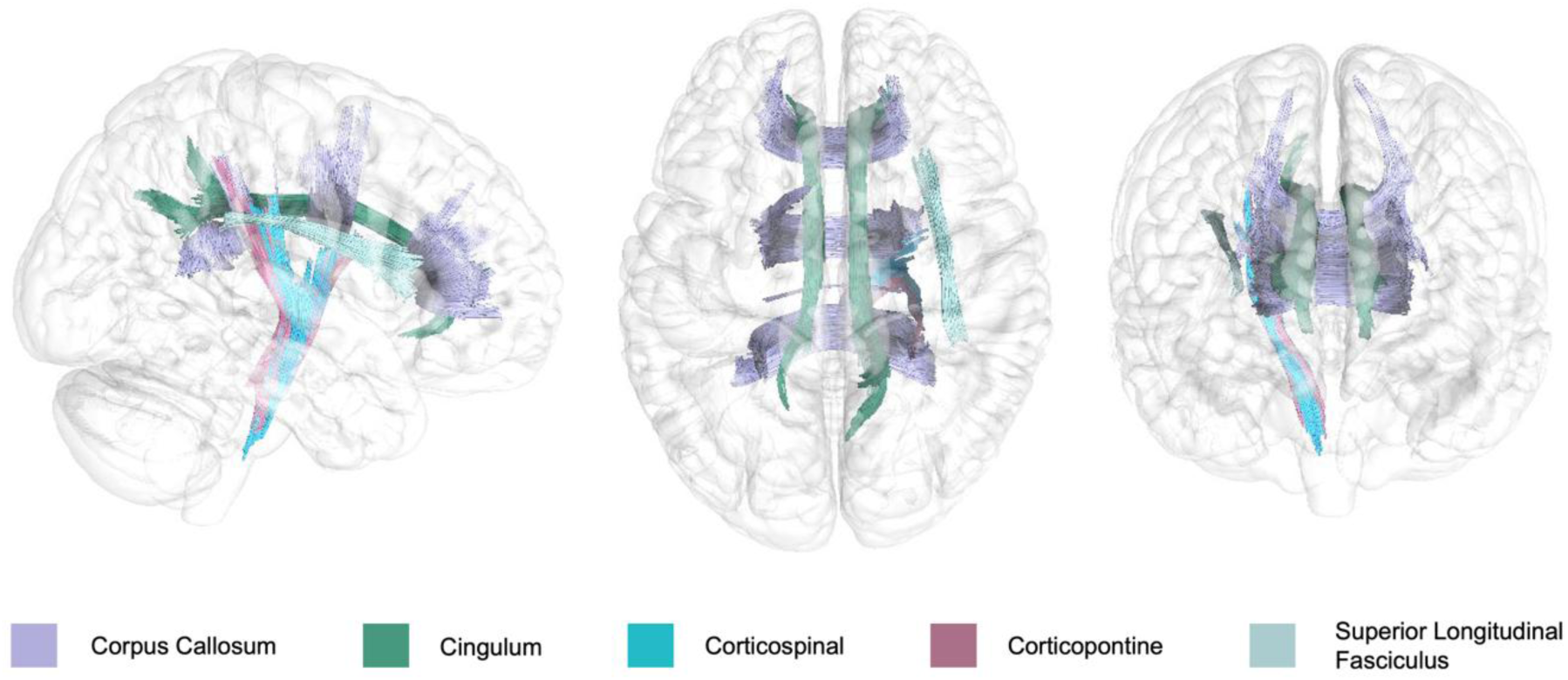
Endometriosis-related pain is associated with white matter microstructure. Correlational tractography in DSI Studio (10,000 permutations, FDR-corrected at *q* < 0.05) revealed positive associations between pain severity and quantitative anisotropy. Top five tracts highlighted for viewing and full tract results are reported in Supplementary Table 8.

## Discussion

Here, we found that endometriosis is associated with higher pain, depression, and anxiety along with specific cognitive deficits, with no observed GMV or white matter microstructure differences at the group level. However, within the endometriosis group, greater pain severity was associated with lower GMV in several temporal and cingulate regions and higher quantitative anisotropy across multiple white matter tracts. Such individual differences within the endometriosis group, despite the absence of case-control differences in GMV and white matter microstructure, suggest that pain and pain-related brain alterations vary along a continuous dimension rather than reflecting categorical group distinctions.

Given the prolonged diagnostic timeline, limited treatment options for chronic pelvic pain, and the debilitating nature of endometriosis, elevated depressive and anxiety symptoms among affected individuals were anticipated. Prior cohort studies and systematic reviews report greater mental health burden in those with endometriosis (6,11) aligning with the present findings. This heightened psychological distress, coupled with persistent pain, may also contribute to the observed baseline differences in cognitive performance (12).The current analyses characterized the negative impact of endometriosis on cognition, and the pattern we observed is consistent with broader chronic pain research demonstrating cognitive impairments, particularly in processing speed, memory, and attention across pain conditions (13). These domains depend on frontoparietal and limbic networks that support both pain modulation and emotional regulation, suggesting overlapping neurocognitive circuitry may underlie the affective and cognitive burden observed here. Supporting this interpretation, a primate model of endometriosis demonstrated impaired long-term memory and learning in marmosets relative to healthy controls (14). Collectively, these findings situate the current work within the broader chronic pain literature while providing the first direct evidence of objective cognitive impairments in humans with endometriosis.

We also examined whether endometriosis-related pain was associated with brain morphology. Greater pain severity was correlated with lower GMV in key pain-related regions, including the left inferior and middle temporal gyri and left cingulate gyrus. This pattern aligns with emerging structural neuroimaging evidence in endometriosis, although prior studies have reported heterogeneous effects. For example, As-Sanie et al. (2012) found reduced GMV in pain-related regions (left cingulate gyrus, thalamus, and right insula) among individuals with endometriosis-associated chronic pelvic pain, with pain ratings negatively correlated with GMV (7). Notably, their small sample size (n < 20 per endometriosis group), reliance on voxel-based morphometry, and lack of intracranial volume control may account for discrepancies with the present findings, where no group-level GMV differences emerged. A more recent investigation by Maulitz and colleagues (2024) addressed several of these methodological limitations and reported increased GMV in the right cerebellum and fusiform gyrus in individuals with endometriosis compared to those without, as well as a negative association between pain severity and cortical thickness in the left temporal lobe (15). This converges with the current study, which similarly found that higher pain was associated with lower GMV in overlapping temporal regions. Jotwani et al. (2025) further identified reduced GMV in the right lateral occipital cortex and left fusiform gyrus areas across a broader age range of individuals with endometriosis compared to controls (8). Together, these studies demonstrate structural brain alterations linked to pain in endometriosis, but differ in the direction and spatial distribution of effects. Such divergence may again reflect methodological variability, particularly differences in analytic resolution, patient population characteristics, sample size, statistical thresholding, and covariate control, underscoring the need for large, harmonized studies to clarify structural brain alterations in endometriosis. Rather than a categorical diagnostic effect, our results suggest that brain morphology varies dimensionally with pain experience.

To our knowledge, the present study represents the first systematic characterization of brain-based white matter microstructure in endometriosis. We found that within the case group, reported endometriosis pain was positively correlated with quantitative anisotropy across interhemispheric, limbic, and frontoparietal pathways implicated in cognitive and affective aspects of pain (corpus callosum, cingulum, superior longitudinal fasciculus) as well as sensorimotor pathways (corticospinal and corticopontine tracts). These associations suggest that endometriosis-related pain engages distributed rather than localized white matter pathways and that white matter organization varies dimensionally with pain severity rather than categorically with diagnosis, in keeping with the GMV findings. Emerging evidence of white matter alterations in chronic pain conditions converge with the present study. Widespread white matter microstructural changes have been reported in patients with dysmenorrhea compared to healthy controls (16), and in relation to dysmenorrhea severity (17). In a large adolescent cohort, pain intensity was associated with white matter microstructure in association tracts proposed to be involved in integrating the pain network (18).These studies demonstrate that chronic pain is linked to specific patterns of white matter reorganization and the current findings extend this framework to endometriosis, highlighting structural alterations in tracts supporting sensory, affective, and cognitive aspects of pain.

While these findings provide new insight into the neurobiology of endometriosis, several caveats should be acknowledged. The case group included comorbidities such as depression, anxiety, and other gynecological disorders. Although this heterogeneity introduces potential confounds, it reflects the clinical reality of endometriosis; excluding such factors would yield an artificially narrow view of the condition. Many participants were using hormonal contraceptives, a common first-line treatment for endometriosis-related pain. However, hormonal contraceptive status was aligned between cases and controls and endocrine assays revealed no group differences in estradiol, progesterone, testosterone, luteinizing hormone (LH), or follicle-stimulating hormone (FSH), suggesting that observed neural or behavioral differences were unlikely to be driven by endogenous hormone levels. Further, cases did not undergo surgical confirmation of endometriosis. While laparoscopy remains the diagnostic standard, there is growing consensus among clinicians and researchers that broader clinical diagnostic criteria should be adopted to reduce diagnostic delays (19). Endometriosis cases in this study were diagnosed at the UCSD Center for Endometriosis Research and Treatment and partnering endometriosis clinics using these published clinical criteria (including evaluation of symptoms, reviewing patient history, and performing physical examinations and/or imaging) by an experienced clinician (coauthor S.A).

The present study advances our understanding of endometriosis-related brain changes by integrating behavioral, cognitive, and multimodal neuroimaging measures in a highly characterized cohort of endometriosis patients and, for the first time, providing a detailed characterization of brain white matter structure. Looking ahead, understanding how hormone-based treatments for endometriosis-related pain influence pain circuitry, including their potential to stall or reverse grey and white matter alterations, represents a critical next step toward translating these findings into clinical care.

The past decade has been a transformative period for understanding how the brain is shaped by female-specific experiences (20–24). This growing body of work has laid a powerful foundation for centering women’s health in neuroimaging. An exciting, much-needed next step is to extend these frameworks to individuals with endocrine disorders, including people with endometriosis (25).This study provides the most comprehensive evidence to date linking pain severity to brain morphology in endometriosis. In doing so, it highlights that reproductive conditions are not anomalies to be excluded but critical sources for painting an accurate, inclusive picture of women’s brain health.

## Methods

### Participants

The current study includes participants recruited as part of an ongoing multicenter longitudinal study at UCSB or UCSD (**Figure 1**). English-speaking women between the ages of 18 to 45 years old were recruited as participants for the study. Participants in the case group (n = 60) were enrolled in the study if they had endometriosis diagnosed by a physician and were intending to begin medication to mitigate endometriosis-related pain. Control group participants (n = 60) were either naturally cycling or on a progestin-only intrauterine device or arm implant, designed to match the endocrine state of the endometriosis participants before beginning medication. Exclusionary criteria for controls included diagnosis of endometriosis, MRI contraindications, history of neurological or endocrine disorders, current pregnancy or breastfeeding, and current use of non-progestin IUD contraceptives.

### Study Design

Prior to the study visit, participants completed a take-home questionnaire including comprehensive assessments of demographics, reproductive health, general health, and mood. Study visits took place at the UCSB Cognitive Neuroendocrinology Laboratory and the UCSB Brain Imaging Center or the UCSD Center for Functional MRI. Visits included comprehensive neuropsychological assessments, a 1.5-hour structural and functional MRI scan, and serological hormone assessments. This study was approved by UCSD’s institutional review board and a UC Reliance with UCSB; all participants provided written informed consent and were compensated for their participation.

### Endocrine Assessments

Participants underwent a blood draw to evaluate hypothalamic-pituitary-gonadal axis hormones, including serum levels of gonadal hormones (17β-estradiol, progesterone, and testosterone) and the pituitary gonadotropins (LH and FSH). One 10 mL blood sample was collected in a vacutainer SST (BD Diagnostic Systems) at each session. The sample clotted at room temperature for 45 min until centrifugation (2000×g for 10 min) and serum was then aliquoted into 1 mL microtubes. Serum samples were stored at −80 °C until assayed. Serum concentrations were determined via liquid chromatography mass-spectrometry (for all steroid hormones) and immunoassay (for all gonadotropins) at the Brigham and Women’s Hospital Research Assay Core. Assay sensitivities, dynamic range, and intra-assay coefficients of variation (respectively) were as follows: estradiol, 1 pg/mL, 1–500 pg/mL, < 5% relative standard deviation (RSD); progesterone, 0.05 ng/mL, 0.05-10 ng/mL, 9.33% RSD; testosterone, 1.0 ng/dL, 1-2000 ng/dL, <4% RSD. FSH and LH levels were determined via chemiluminescent assay (Beckman Coulter). The assay sensitivity, dynamic range, and the intra-assay coefficient of variation were as follows: FSH, 0.2 mIU/mL, 0.2-200 mIU/mL, 3.1-4.3%; LH, 0.2 mIU/mL, 0.2-250 mIU/mL, 4.3-6.4%. Progesterone values < 0.05 were set at 0.05. LH and FSH levels < 0.2 were set to 0.2.

### Questionnaires

The following scales were administered as part of the take-home questionnaires: Adverse Childhood Experience Questionnaire (26), Endometriosis Health Profile (27), Center for Epidemiologic Studies Depression Scale (28), State-Trait Anxiety Inventory for Adults (29), Biberoglu and Berman Scale (30), Medical Outcomes Study Short Form (31), Pittsburgh Sleep Quality Index (32), International Physical Activity (33), the Post-Traumatic Stress Disorder Checklist-Civilian Version (34), and the Brief Pain Inventory Short Form (35).

### Cognitive Assessments

Participants completed a neuropsychological battery including the following assessments: Selective Reminding Task (36) Digit Span (37) Digit Symbol Coding (38) Stroop (39), and Delis-Kaplan Executive Function System: Verbal Fluency (40). They also completed the Face-Name Memory Exam, an assessment of associative memory that has been shown to be impacted by periods of hormonal fluctuation (41, 42) and a curated cognitive battery (Oral Symbol Digit Test, Picture Vocabulary Test, Oral Reading Recognition, Picture Sequence Memory Test, List Sorting, Pattern Comparison, Flanker, Dimensional Change Card Sort) from the well-validated NIH Toolbox (43).

### MRI Acquisition

Participants underwent a whole-brain MRI on a Siemens 3 T Prisma scanner equipped with a 64-channel phased-array head coil. High-resolution anatomical scans were acquired using a T1-weighted magnetization-prepared rapid gradient echo (MPRAGE) sequence (TR, 2,500 ms; TE, 2.31 ms; TI, 934 ms; flip angle, 7°; 0.8 mm thickness), followed by a gradient echo fieldmap (TR, 758 ms; TE1, 4.92 ms; TE2, 7.38 ms; flip angle, 60°). A Diffusion Spectrum Imaging (DSI) scan was also acquired with the following parameters: TR, 4300 ms, echo time, 100.20 ms, 139 directions, b-max, 5,000, FoV, 259 mm, 87 slices, 1.8 *x* 1.8 *x* 1.8 mm voxel resolution.

### Brain morphology assessments

Measures of cerebrospinal fluid, ventricle size, cortical GMV and cortical thickness and subcortical volumes were calculated by running T1w images through the recon-all pipeline from FreeSurfer (44, 45). Cortical measures were segmented into the 68-region Desikan-Killiany atlas and subcortical measures were segmented into 28 regions-of-interest determined via the aseg parcellation (46). Global morphology and GMV measures were corrected for total intracranial volume.

### White Matter Microstructure

Diffusion-weighted images were processed using QSIPrep (version 0.23.2) with default parameters, except for the following modifications: unringing method = mridegibbs, HMC model = 3dSHORE, b0-to-T1 transform = Affine, intramodal template transform = SyN, and output resolution = 1.8 mm. Quality assurance was conducted using the QSIPrep-generated html reports and desc-image_qc.tsv files before proceeding to reconstruction. Reconstruction was performed with QSIRecon’s GQI plus deterministic tractography pipeline (dsi_studio_gqi). We assessed differences in global white matter microstructure (e.g., average whole-brain quantitative anisotropy). Here, we took the voxelwise scalar maps returned by the GQI pipeline (in subject T1w space) and warped them into MNI152NLin2009cAsym space using the composite transform derived via QSIPrep. We then computed a weighted mean across the whole brain given a white matter probability map in the MNI template space. The relationship between endometriosis-specific pain and white matter microstructure (quantitative anisotropy) was then examined using Correlational Tractography in DSI Studio. The following parameters were applied: FDR threshold *p* < 0.05, pruning iterations = 16, permutation count = 10,000, t-score threshold = 2.5, and minimum tract length = 25 voxels. Only regions containing at least 10 tracts are reported.

### Statistical Analyses

All statistical analyses were conducted in R (version 4.3.2). Analyses were two-tailed where applicable. To assess mood, cognitive, and neural differences between the control and endometriosis groups, multivariate regressions (FDR-corrected) and independent samples t-tests (Bonferroni-corrected) were performed. For variables that did not meet normality assumptions (BMI, hormones), Wilcoxon rank-sum tests were used. Fisher’s exact tests were conducted to compare race and ethnicity across groups. Pearson’s correlations and partial correlations were computed between brain, pain, and cognitive measures (FDR-corrected).

## Acknowledgements

We thank M. Mendoza (UCSB) for assistance with scanning and phlebotomy, the USC Alzheimer’s Therapeutic Research Institute for blood sample storage and processing and A. Jackson of the UCSD Center for Function MRI for assistance with imaging procedures. Finally, we thank the many undergraduate research assistants whose efforts made data collection possible.

## Data Availability Statement

The data that support the findings of this study are part of a larger, ongoing project involving a sensitive patient population. As such, the data are not publicly available at this time. No custom code was used.

