## Supplementary Material for "Neurobiological Signatures of Endometriosis: Characterizing Pain, Cognition, and Brain Morphology"

| Birth Control Type | Control (n = 60) | Case (n = 60) |
|---|---|---|
| **None** | 35 (58.33%) | 37 (61.67%) |
| **Intrauterine Device** | 21 (35.00%) | 10 (16.67%) |
| **Arm Implant** | 4 (6.67%) | 4 (6.67%) |
| **Combination Pill** | NA | 5 (8.33%) |
| **Multiple** | NA | 1 (1.67%) |
| **Progesterone Pill** | NA | 1 (1.67%) |
| **Vaginal Ring** | NA | 2 (3.33%) |

**Supplementary Table 1. Birth Control Type.**

| Questionnaire Assessment | **Control** Mean (SD) \| Range | **Case** Mean (SD) \| Range | p-value |
|---|---|---|---|
| **Migraine** | **0.63 (1.31) \| 0-5** | **2.93 (2.57) \| 0-6** | **1.79E-07** |
| **Physical Function** | **96.25 (9.59) \| 35-100** | **75.73 (24.16) \| 10-100** | **2.68E-07** |
| **Physical Limit** | **91.95 (23.42) \| 0-100** | **43.64 (42.82) \| 0-100** | **4.45E-10** |
| **Emotional Limit** | **67.78 (38.8) \| 0-100** | **49.7 (42.49) \| 0-100** | **2.04E-02** |
| **Energy Fatigue** | **55 (20.91) \| 0-95** | **32.27 (20.45) \| 0-85** | **1.16E-07** |
| **Emotion Well-Being** | **68.07 (18.09) \| 12-100** | **58.76 (18.73) \| 16-88** | **8.97E-03** |
| **Social Function** | **82.71 (20.85) \| 0-100** | **57.05 (24.97) \| 0-100** | **1.16E-07** |
| **Pain Score** | **83.12 (16.71) \| 22.5-100** | **36.09 (21.59) \| 0-77.5** | **2.28E-22** |
| **General Health** | **75.67 (13.23) \| 30-100** | **48 (20.52) \| 5-90** | **1.90E-12** |
| **Total ACE** | **1.54 (1.60) \| 0-7** | **3.29 (2.64) \| 0-9** | **8.42E-05** |
| **Total MET** | 2964.85 (2311.07) \| 198-13356 | 3540.11 (3975.07) \| 0-19572 | 3.54E-01 |
| **Total STAI** | **34.88 (10.63) \| 20-56** | **41.69 (11.98) \| 20-73** | **2.26E-03** |
| **Total CESD** | **8.67 (8.26) \| 0-39** | **16.76 (11.07) \| 0-43** | **4.99E-05** |
| **PTSD Score** | **28.33 (11.04) \| 17-77** | **36.11 (13.79) \| 18-74** | **1.75E-03** |
| **Total Pelvic Pain** | **1.47 (1.32) \| 0-5** | **6.3 (1.62) \| 2-9** | **7.48E-28** |
| **Chronic Pain** | **1.06 (1.57) \| 0-5.14** | **4.85 (2.36) \| 0.29-10** | **9.83E-06** |
| **PSQI** | **5.13 (3.11) \| 0-12** | **7.59 (3.52) \| 2-16** | **2.09E-04** |

**Supplementary Table 2. Reported Mood and Lifestyle Scores.** All FDR-corrected at $q < 0.05$. ACE = Adverse Childhood Events, MET = Metabolic Equivalent Time/ week, STAI = State-Trait Anxiety Inventory, CES-D = Center for Epidemiologic Studies Depression scale , PTSD = Post-Traumatic Stress Disorder, PSQI = Pittsburgh Sleep Quality Index.

| Cognitive Assessment | Control<br>Mean (SD) \| Range | Case<br>Mean (SD) \| Range | p-value |
|---|---|---|---|
| **SRT: Total Recall** | 57.70 (5.68) \| 44-68 | 53.43 (8.58) \| 32-69 | 0.005 |
| **SRT: Long-Term Recall** | 53.03 (8.45) \| 35-68 | 47.29 (12.41) \| 15-68 | 0.009 |
| **SRT: Long-Term Storage** | 54.40 (7.97) \| 40-68 | 50.14 (11.93) \| 16-68 | 0.04 |
| **SRT: Consistent Long-Term Retrieval** | 48.45 (11.50) \| 17-68 | 40.60 (14.33) \| 10-68 | 0.004* |
| **Digit Span Forward: Max** | 7.10 (1.51) \| 4-11 | 6.81 (1.26) \| 4-10 | 0.28 |
| **Digit Span Backward: Max** | 5.48 (1.19) \| 3-9 | 4.86 (1.15) \| 3-8 | 0.009* |
| **Stroop: Words** | 96.78 (5.56) \| 80-100 | 92.29 (9.99) \| 63-100 | 0.008* |
| **Stroop: Colors** | 76.33 (10.60) \| 54-100 | 71.84 (11.38) \| 48-95 | 0.04* |
| **Stroop: Colors and Words** | 51.53 (10.89) \| 28-80 | 46.72 (10.76) \| 24-72 | 0.03* |
| **SRT: Delayed Recall** | 10.57 (1.45) \| 6-12 | 9.43 (2.10) \| 3-12 | 0.004 |
| **SRT: Delayed Recognition** | 11.97 (0.18) \| 11-12 | 11.91 (0.34) \| 10-12 | 0.30 |
| **Fluencies: Category Words** | 16.03 (2.64) \| 8-21 | 15.31 (2.41) \| 9-20 | 0.15 |
| **Fluencies: Category Switching** | 14.67 (2.78) \| 6-20 | 13.97 (2.62) \| 7-20 | 0.18 |
| **FNAME: Initial Recall Names** | 14.32 (3.85) \| 6-22 | 11.38 (4.33) \| 3-21 | 0.002* |
| **FNAME: Initial Recall Occupations** | 18.88 (3.39) \| 10-24 | 17.39 (4.23) \| 4-24 | 0.05 |
| **FNAME: Initial Recall Total** | 33.2 (6.45) \| 19-45 | 28.77 (7.77) \| 8-44 | 0.004* |
| **FNAME: Names Total** | 24.02 (6.10) \| 12-34 | 19.09 (7.30) \| 4-33 | 0.002* |
| **FNAME: Occupations Total** | 29.92 (4.69) \| 17-36 | 27.70 (6.10) \| 7-36 | 0.04 |
| **FNAME: Total** | 53.93 (9.90) \| 31-69 | 46.79 (12.42) \| 12-68 | 0.004* |

**Supplementary Table 3. Neuropsychological Battery.** All FDR-corrected at $q < 0.05$. SRT = Selective Reminding Task, FNAME = Face-Name Associative Memory Exam. *Endometriosis group remained a significant negative predictor of cognitive performance after controlling for years of education.

| Cognitive Assessment | Control<br>Mean (SD) \| Range | Case<br>Mean (SD) \| Range | p-value |
|---|---|---|---|
| **Oral Symbol Digit Coding: Raw Score** | 100.3 (15.47) \| 68-139 | 94.16 (13.50) \| 62-132 | 0.03 |
| **List Sorting: Raw Score** | 19.3 (3.00) \| 14-26 | 18.33 (2.52) \| 12-25 | 0.07 |
| **Pattern Comparison: Raw Score** | 54.72 (9.20) \| 37-76 | 49.31 (9.32) \| 27-67 | 0.005* |
| **Picture Vocabulary: Theta Score** | 5.85 (1.89) \| 1.73-10.63 | 4.7 (1.84) \| 1.05-8.63 | 0.004 |
| **Oral Reading Recognition: Theta Score** | 7.07 (2.02) \| 1.98-10.3 | 5.73 (2.48) \| -1.17-9.92 | 0.005 |
| **Picture Sequence: Theta Score** | 0.46 (0.79) \| -1.83-1.55 | 0.08 (0.81) \| -1.62-1.55 | 0.02 |
| **Picture Sequence: Computed Score** | 630.93 (78.85) \| 402.14-740.11 | 593.24 (81.19) \| 422.54-740.11 | 0.02 |
| **Flanker: Computed Score** | 8.82 (0.67) \| 7.33-10 | 8.61 (0.71) \| 6.44-10 | 0.09 |
| **DCCST: Computed Score** | 9.01 (0.72) \| 7.3-10 | 8.47 (1.30) \| 3.75-10 | 0.01 |
| **Fluid Cognition Composite Score** | 111.33 (16.15) \| 73-144 | 101.96 (16.6) \| 65-138 | 0.005† |
| **Crystallized Cognition Composite Score** | 117.9 (14.00) \| 80-146 | 106.83 (15.23) \| 71-137 | 0.0005*† |
| **Cognitive Function Composite Score** | 116.92 (14.10) \| 73-146 | 105.28 (14.18) \| 74-135 | 0.0002*† |

**Supplementary Table 4. NIH Toolbox Measures.** All FDR-corrected at $q < 0.05$. DCCST = Dimensional Change Card Sort Task. *Endometriosis group remained a significant negative predictor of cognitive performance after controlling for years of education. †NIH Toolbox-derived age-corrected standard score.

| Brain Region | Control<br>Mean (SD) \| Range | Case<br>Mean (SD) \| Range | p-value |
|---|---|---|---|
| **Cortex Volume** | 0.3658 (0.0427) \| 0.2859-0.4409 | 0.3765 (0.0404) \| 0.2997-0.4689 | 0.39 |
| **Total GMV** | 0.4931 (0.0581) \| 0.3882-0.607 | 0.51 (0.0533) \| 0.4078-0.6175 | 0.37 |
| **Subcortical GMV** | 0.0465 (0.0056) \| 0.0366-0.0598 | 0.0483 (0.0056) \| 0.0386-0.0601 | 0.34 |
| **Cerebral WMV** | 0.3518 (0.0413) \| 0.2852-0.4554 | 0.3593 (0.0415) \| 0.2933-0.4453 | 0.53 |
| **Cerebrospinal Fluid** | 7e-04 (1e-04) \| 5e-04-0.0012 | 7e-04 (2e-04) \| 4e-04-0.0011 | 0.39 |
| **Left Lateral Ventricle** | 0.0049 (0.0019) \| 0.0019-0.0131 | 0.0049 (0.0018) \| 0.0024-0.0103 | 0.90 |
| **Left Inferior Lateral Ventricle** | 2e-04 (1e-04) \| 1e-04-6e-04 | 3e-04 (1e-04) \| 1e-04-8e-04 | 0.59 |
| **Left Cerebellum WM** | 0.0109 (0.0015) \| 0.0088-0.0142 | 0.0114 (0.0016) \| 0.0085-0.0159 | 0.39 |
| **Left Cerebellum Cortex** | 0.04 (0.0059) \| 0.0306-0.0565 | 0.0422 (0.0054) \| 0.0321-0.052 | 0.34 |
| **Left Thalamus** | 0.0058 (8e-04) \| 0.0045-0.0074 | 0.0061 (9e-04) \| 0.0045-0.0092 | 0.34 |
| **Left Caudate** | 0.0028 (4e-04) \| 0.0019-0.0038 | 0.0029 (4e-04) \| 0.0021-0.0039 | 0.59 |
| **Left Putamen** | 0.004 (5e-04) \| 0.0031-0.0059 | 0.0042 (6e-04) \| 0.0032-0.0055 | 0.34 |
| **Left Pallidum** | 0.0016 (2e-04) \| 0.0013-0.002 | 0.0017 (2e-04) \| 0.0013-0.0023 | 0.34 |
| **Third Ventricle** | 7e-04 (2e-04) \| 4e-04-0.0013 | 7e-04 (1e-04) \| 5e-04-0.0012 | 0.53 |
| **Fourth Ventricle** | 0.0013 (4e-04) \| 7e-04-0.0023 | 0.0012 (3e-04) \| 7e-04-0.0021 | 0.53 |
| **Brainstem** | 0.0161 (0.0024) \| 0.0121-0.0216 | 0.0168 (0.0023) \| 0.0128-0.0223 | 0.34 |
| **Left Hippocampus** | 0.0032 (4e-04) \| 0.0025-0.0045 | 0.0034 (4e-04) \| 0.0026-0.0047 | 0.34 |
| **Left Amygdala** | 0.0014 (3e-04) \| 0.001-0.0022 | 0.0014 (2e-04) \| 0.001-0.002 | 0.38 |
| **Left Accumbens** | 5e-04 (1e-04) \| 3e-04-9e-04 | 5e-04 (1e-04) \| 4e-04-7e-04 | 0.59 |
| **Left Ventral Diencephalon** | 0.0031 (4e-04) \| 0.0024-0.0041 | 0.0032 (4e-04) \| 0.0025-0.0043 | 0.39 |
| **Left Choroid** | 4e-04 (1e-04) \| 2e-04-6e-04 | 4e-04 (1e-04) \| 2e-04-7e-04 | 0.59 |
| **Right Lateral Ventricle** | 0.0045 (0.0018) \| 0.0016-0.0136 | 0.0043 (0.0017) \| 0.0019-0.0113 | 0.67 |
| **Right Inferior Lateral Ventricle** | 3e-04 (1e-04) \| 1e-04-7e-04 | 3e-04 (1e-04) \| 1e-04-5e-04 | 0.85 |
| **Right Cerebellum WM** | 0.0105 (0.0015) \| 0.0085-0.0147 | 0.0109 (0.0015) \| 0.0082-0.015 | 0.39 |
| **Right Cerebellum Cortex** | 0.0409 (0.0062) \| 0.0314-0.057 | 0.0432 (0.0053) \| 0.0332-0.0531 | 0.34 |
| **Right Thalamus** | 0.0058 (8e-04) \| 0.0045-0.0076 | 0.006 (8e-04) \| 0.0046-0.0084 | 0.39 |
| **Right Caudate** | 0.003 (4e-04) \| 0.0019-0.004 | 0.003 (4e-04) \| 0.0022-0.0039 | 0.71 |
| **Right Putamen** | 0.0041 (6e-04) \| 0.0032-0.0062 | 0.0043 (6e-04) \| 0.0033-0.0055 | 0.38 |
| **Right Pallidum** | 0.0015 (2e-04) \| 0.0011-0.0019 | 0.0015 (2e-04) \| 0.0012-0.002 | 0.39 |
| **Right Hippocampus** | 0.0033 (4e-04) \| 0.0025-0.0046 | 0.0034 (4e-04) \| 0.0027-0.0049 | 0.34 |
| **Right Amygdala** | 0.0013 (2e-04) \| 9e-04-0.0018 | 0.0013 (2e-04) \| 0.001-0.0017 | 0.67 |
| **Right Accumbens** | 5e-04 (1e-04) \| 4e-04-8e-04 | 5e-04 (1e-04) \| 3e-04-8e-04 | 0.67 |
| **Right Ventral Diencephalon** | 0.0032 (4e-04) \| 0.0025-0.0041 | 0.0033 (4e-04) \| 0.0025-0.0044 | 0.53 |
| **Right Choroid** | 5e-04 (1e-04) \| 3e-04-8e-04 | 5e-04 (1e-04) \| 3e-04-9e-04 | 0.39 |
| **Optic Chiasm** | 1e-04 (0) \| 1e-04-2e-04 | 1e-04 (0) \| 1e-04-2e-04 | 0.55 |

**Supplementary Table 5. Brain Structural Measures.** All FDR-corrected at $q < 0.05$. GMV = Gray Matter Volume. WMV = White Matter Volume.

| Brain Region | Control<br>Mean (SD) \| Range | Case<br>Mean (SD) \| Range | p-value |
|---|---|---|---|
| **lh_bankssts** | 0.0019 (3e-04) \| 0.0011-0.0025 | 0.0018 (4e-04) \| 0.0011-0.0027 | 0.84 |
| **lh_caudalanteriorcingulate** | 0.0014 (3e-04) \| 5e-04-0.0023 | 0.0014 (4e-04) \| 8e-04-0.0027 | 0.92 |
| **lh_caudalmiddlefrontal** | 0.0048 (9e-04) \| 0.0032-0.0069 | 0.0047 (7e-04) \| 0.0028-0.0067 | 0.98 |
| **lh_cuneus** | 0.0024 (4e-04) \| 0.0017-0.0036 | 0.0026 (6e-04) \| 0.0017-0.0042 | 0.42 |
| **lh_entorhinal** | 0.0014 (2e-04) \| 9e-04-0.002 | 0.0015 (3e-04) \| 0.0011-0.0023 | 0.43 |
| **lh_fusiform** | 0.0073 (0.001) \| 0.0048-0.0092 | 0.0077 (0.0011) \| 0.0057-0.0101 | 0.31 |
| **lh_inferiorparietal** | 0.0095 (0.0015) \| 0.007-0.0131 | 0.0098 (0.0013) \| 0.0077-0.013 | 0.50 |
| **lh_inferiortemporal** | 0.0084 (0.0015) \| 0.0056-0.0117 | 0.0087 (0.0013) \| 0.0057-0.0113 | 0.50 |
| **lh_isthmuscingulate** | 0.0021 (3e-04) \| 0.0015-0.0029 | 0.0022 (3e-04) \| 0.0013-0.0029 | 0.63 |
| **lh_lateraloccipital** | 0.0096 (0.0015) \| 0.0069-0.0128 | 0.0101 (0.0017) \| 0.0073-0.0137 | 0.42 |
| **lh_lateralorbitofrontal** | 0.0058 (7e-04) \| 0.0045-0.0076 | 0.006 (7e-04) \| 0.0047-0.0078 | 0.48 |
| **lh_lingual** | 0.0052 (0.001) \| 0.0035-0.0076 | 0.0055 (9e-04) \| 0.0035-0.0075 | 0.42 |
| **lh_medialorbitofrontal** | 0.0037 (5e-04) \| 0.0026-0.0049 | 0.0037 (5e-04) \| 0.0029-0.0047 | 0.98 |
| **lh_middletemporal** | 0.0091 (0.0013) \| 0.0067-0.012 | 0.0091 (0.0014) \| 0.0064-0.012 | 0.93 |
| **lh_parahippocampal** | 0.0017 (3e-04) \| 0.0012-0.0024 | 0.0018 (3e-04) \| 0.0012-0.0026 | 0.47 |
| **lh_paracentral** | 0.0028 (5e-04) \| 0.0018-0.0037 | 0.0029 (4e-04) \| 0.0022-0.0039 | 0.42 |
| **lh_parsopercularis** | 0.0037 (8e-04) \| 0.0022-0.0071 | 0.0039 (7e-04) \| 0.0027-0.0059 | 0.50 |
| **lh_parsorbitalis** | 0.0018 (3e-04) \| 0.0013-0.0026 | 0.002 (3e-04) \| 0.0012-0.0027 | 0.31 |
| **lh_parstriangularis** | 0.003 (5e-04) \| 0.0019-0.0042 | 0.0031 (6e-04) \| 0.0021-0.0047 | 0.48 |
| **lh_pericalcarine** | 0.0016 (4e-04) \| 0.001-0.0027 | 0.0018 (5e-04) \| 9e-04-0.0034 | 0.37 |
| **lh_postcentral** | 0.0074 (0.001) \| 0.0056-0.0101 | 0.0077 (0.0011) \| 0.0055-0.0102 | 0.43 |
| **lh_posteriorcingulate** | 0.0025 (4e-04) \| 0.0014-0.0034 | 0.0025 (4e-04) \| 0.0018-0.0034 | 0.99 |
| **lh_precentral** | 0.0106 (0.0015) \| 0.0076-0.014 | 0.011 (0.0014) \| 0.0084-0.0142 | 0.42 |
| **lh_precuneus** | 0.0075 (0.0011) \| 0.0057-0.01 | 0.0076 (0.001) \| 0.006-0.0095 | 0.92 |
| **lh_rostralanteriorcingulate** | 0.002 (4e-04) \| 0.0014-0.0029 | 0.0019 (4e-04) \| 0.001-0.0029 | 0.50 |
| **lh_rostralmiddlefrontal** | 0.0115 (0.0019) \| 0.0079-0.0168 | 0.0116 (0.0017) \| 0.0079-0.0152 | 0.92 |
| **lh_superiorfrontal** | 0.0167 (0.0024) \| 0.0118-0.0215 | 0.0172 (0.002) \| 0.0124-0.0215 | 0.43 |
| **lh_superiorparietal** | 0.0101 (0.0017) \| 0.0068-0.0144 | 0.0106 (0.0019) \| 0.0068-0.0165 | 0.43 |
| **lh_superiortemporal** | 0.0101 (0.0014) \| 0.0076-0.0135 | 0.0104 (0.0016) \| 0.0074-0.0139 | 0.50 |
| **lh_supramarginal** | 0.0089 (0.0015) \| 0.0048-0.0127 | 0.009 (0.0014) \| 0.0068-0.0123 | 0.98 |
| **lh_frontalpole** | 7e-04 (1e-04) \| 5e-04-0.0011 | 8e-04 (1e-04) \| 5e-04-0.0011 | 0.87 |
| **lh_temporalpole** | 0.0019 (3e-04) \| 0.0011-0.0029 | 0.0019 (3e-04) \| 0.0011-0.0029 | 0.50 |
| **lh_transversetemporal** | 0.001 (2e-04) \| 6e-04-0.0015 | 0.001 (3e-04) \| 5e-04-0.0022 | 0.43 |
| **lh_insula** | 0.0053 (7e-04) \| 0.0039-0.0069 | 0.0055 (7e-04) \| 0.0042-0.0069 | 0.43 |
| **rh_bankssts** | 0.0017 (3e-04) \| 0.0011-0.0024 | 0.0018 (3e-04) \| 0.0012-0.0027 | 0.83 |
| **rh_caudalanteriorcingulate** | 0.0016 (4e-04) \| 8e-04-0.0025 | 0.0016 (3e-04) \| 6e-04-0.0023 | 0.84 |
| **rh_caudalmiddlefrontal** | 0.0044 (9e-04) \| 0.0029-0.0079 | 0.0043 (9e-04) \| 0.0026-0.0067 | 0.51 |
| **rh_cuneus** | 0.0027 (5e-04) \| 0.0016-0.0041 | 0.0029 (6e-04) \| 0.0014-0.0047 | 0.42 |
| **rh_entorhinal** | 0.0015 (3e-04) \| 9e-04-0.0021 | 0.0017 (3e-04) \| 0.0011-0.0024 | 0.31 |
| **rh_fusiform** | 0.0073 (0.0012) \| 0.005-0.0101 | 0.0077 (0.0011) \| 0.0057-0.0104 | 0.42 |
| **rh_inferiorparietal** | 0.0116 (0.0018) \| 0.0077-0.0158 | 0.012 (0.0017) \| 0.0091-0.0158 | 0.48 |
| **rh_inferiortemporal** | 0.0081 (0.0014) \| 0.0058-0.0112 | 0.0084 (0.0012) \| 0.0063-0.0109 | 0.43 |
| **rh_isthmuscingulate** | 0.0019 (4e-04) \| 0.0013-0.0028 | 0.002 (3e-04) \| 0.0013-0.0028 | 0.47 |
| **rh_lateraloccipital** | 0.0099 (0.0016) \| 0.0066-0.0153 | 0.0103 (0.0019) \| 0.0065-0.0152 | 0.43 |
| **rh_lateralorbitofrontal** | 0.0054 (7e-04) \| 0.0039-0.0069 | 0.0054 (8e-04) \| 0.004-0.0071 | 0.89 |
| **rh_lingual** | 0.0056 (0.001) \| 0.004-0.008 | 0.006 (0.001) \| 0.0042-0.0083 | 0.31 |
| **rh_medialorbitofrontal** | 0.0039 (6e-04) \| 0.0028-0.0058 | 0.0039 (5e-04) \| 0.0027-0.0049 | 0.99 |
| **rh_middletemporal** | 0.0095 (0.0014) \| 0.0072-0.0131 | 0.0098 (0.0012) \| 0.0079-0.013 | 0.50 |
| **rh_parahippocampal** | 0.0016 (3e-04) \| 0.0011-0.0033 | 0.0016 (3e-04) \| 0.0012-0.0024 | 0.63 |
| **rh_paracentral** | 0.0031 (5e-04) \| 0.002-0.0044 | 0.0032 (4e-04) \| 0.0021-0.004 | 0.50 |
| **rh_parsopercularis** | 0.003 (5e-04) \| 0.0021-0.0042 | 0.0031 (5e-04) \| 0.0022-0.0043 | 0.48 |
| **rh_parsorbitalis** | 0.0022 (4e-04) \| 0.0014-0.0031 | 0.0022 (4e-04) \| 0.0016-0.0033 | 0.50 |
| **rh_parstriangularis** | 0.0033 (6e-04) \| 0.0022-0.0056 | 0.0035 (8e-04) \| 0.0021-0.0055 | 0.43 |
| **rh_pericalcarine** | 0.0018 (4e-04) \| 0.0012-0.0028 | 0.002 (5e-04) \| 0.0012-0.0035 | 0.31 |
| **rh_postcentral** | 0.007 (0.001) \| 0.0048-0.0096 | 0.0073 (0.001) \| 0.0055-0.0098 | 0.43 |
| **rh_posteriorcingulate** | 0.0025 (4e-04) \| 0.0016-0.0037 | 0.0025 (4e-04) \| 0.0018-0.0033 | 0.98 |
| **rh_precentral** | 0.0099 (0.0015) \| 0.0061-0.013 | 0.0102 (0.0015) \| 0.0057-0.0126 | 0.46 |
| **rh_precuneus** | 0.0079 (0.0012) \| 0.0061-0.0114 | 0.0079 (0.0012) \| 0.0053-0.0109 | 0.93 |
| **rh_rostralanteriorcingulate** | 0.0014 (3e-04) \| 6e-04-0.0019 | 0.0015 (3e-04) \| 8e-04-0.002 | 0.47 |
| **rh_rostralmiddlefrontal** | 0.0116 (0.0022) \| 0.008-0.0189 | 0.0116 (0.0021) \| 0.0079-0.0174 | 0.98 |
| **rh_superiorfrontal** | 0.0159 (0.0024) \| 0.011-0.0211 | 0.016 (0.0021) \| 0.0115-0.0208 | 0.92 |
| **rh_superiorparietal** | 0.0099 (0.0016) \| 0.0059-0.0134 | 0.0102 (0.0019) \| 0.0063-0.0142 | 0.61 |
| **rh_superiortemporal** | 0.0093 (0.0014) \| 0.0065-0.0124 | 0.0097 (0.0015) \| 0.0066-0.013 | 0.43 |
| **rh_supramarginal** | 0.0077 (0.0013) \| 0.0047-0.0102 | 0.0079 (0.0015) \| 0.0055-0.012 | 0.63 |
| **rh_frontalpole** | 9e-04 (2e-04) \| 6e-04-0.0017 | 9e-04 (2e-04) \| 6e-04-0.0015 | 0.83 |
| **rh_temporalpole** | 0.0021 (4e-04) \| 0.0013-0.003 | 0.0022 (4e-04) \| 0.0016-0.0032 | 0.37 |
| **rh_transversetemporal** | 7e-04 (1e-04) \| 5e-04-0.0011 | 8e-04 (1e-04) \| 5e-04-0.0012 | 0.63 |
| **rh_insula** | 0.0052 (7e-04) \| 0.0039-0.0069 | 0.0055 (8e-04) \| 0.0042-0.0072 | 0.42 |

**Supplementary Table 6. Cortical Gray Matter Volume Measures**. All FDR-corrected at $q < 0.05$. LH = Left Hemisphere, RH = Right Hemisphere.

| White Matter Microstructure Measure | Control<br>Mean (SD) \| Range | Case<br>Mean (SD) \| Range | p-value |
|---|---|---|---|
| Quantitative Anisotropy | 0.1775 (0.0206) \| 0.1117-0.2128 | 0.1773 (0.0185) \| 0.1342-0.2296 | 0.95 |
| Generalized Fractional Anisotropy | 0.0737 (0.0023) \| 0.0684-0.079 | 0.0728 (0.0025) \| 0.066-0.0786 | 0.27 |
| Axial Diffusivity | 1.0597 (0.0149) \| 1.0051-1.0923 | 1.0567 (0.0168) \| 1.0226-1.0943 | 0.81 |
| Mean Diffusivity | 0.777 (0.0146) \| 0.7285-0.8082 | 0.775 (0.0174) \| 0.7486-0.8176 | 0.85 |
| Radial Diffusivity | 0.6356 (0.0154) \| 0.5902-0.6677 | 0.6342 (0.0186) \| 0.6-0.6867 | 0.85 |

**Supplementary Table 7. Summary White Matter Microstructure Measures.** All FDR-corrected at $q < 0.05$.

| Tract Name | Tract Number | Mean Length (mm) | Span (mm) | Total Volume ($mm^3$) |
|---|---|---|---|---|
| Commissure_CorpusCallosum_Body | 1417 | 60.1838 | 21.8124 | 9180 |
| Commissure_CorpusCallosum_ForcepsMinor | 1280 | 56.0283 | 20.8591 | 9349 |
| Association_CingulumR_Parolfactory | 1231 | 66.8969 | 31.5702 | 2964 |
| Commissure_CorpusCallosum_Tapetum | 805 | 60.6563 | 26.0765 | 4890 |
| Association_CingulumR_FrontalParietal | 769 | 63.7537 | 29.8644 | 3391 |
| Association_CingulumL_Parolfactory | 559 | 61.2004 | 28.5375 | 2445 |
| ProjectionBrainstem_CorticospinalTractR | 405 | 57.8602 | 28.0616 | 3523 |
| ProjectionBrainstem_CorticopontineTractR_Parietal | 325 | 59.4846 | 27.7589 | 1849 |
| Association_SuperiorLongitudinalFasciculusR_2 | 157 | 57.72 | 28.1212 | 1239 |
| Association_InferiorFrontoOccipitalFasciculusR | 123 | 53.2259 | 25.5967 | 877 |
| Association_CingulumL_FrontalParietal | 85 | 60.1493 | 28.8099 | 1225 |
| Association_ArcuateFasciculusL | 80 | 54.6603 | 25.0203 | 609 |
| Association_CingulumL_FrontalParahippocampal | 74 | 62.3577 | 30.0463 | 1014 |
| ProjectionBrainstem_CorticobulbarTractR | 74 | 57.2508 | 28.0496 | 1292 |
| Commissure_CorpusCallosum_ForcepsMajor | 61 | 53.8751 | 23.4129 | 1096 |
| ProjectionBrainstem_CorticopontineTractR_Frontal | 53 | 55.9304 | 27.4143 | 836 |
| Association_ArcuateFasciculusR | 21 | 60.201 | 28.3461 | 543 |

**Supplementary Table 7. Quantitative anisotropy tracts positively associated with endometriosis pain.** All FDR-corrected at $q < 0.05$. R = Right hemisphere, L = Left hemisphere.
